# *De novo* chromosome-length assembly of the mule deer (*Odocoileus hemionus*) genome

**DOI:** 10.1101/2021.08.12.456132

**Authors:** Sydney Lamb, Adam M. Taylor, Tabitha A. Hughes, Brock R. McMillan, Randy T. Larsen, Ruqayya Khan, David Weisz, Olga Dudchenko, Erez Lieberman Aiden, Paul B. Frandsen

## Abstract

The mule deer (*Odocoileus hemionus*) is an ungulate species that ranges from western Canada to central Mexico. Mule deer are an essential source of food for many predators, are relatively abundant, and commonly make broad migration movements. A clearer understanding of the mule deer genome can help facilitate knowledge of its population genetics, movements, and demographic history, aiding in conservation efforts. While mule deer are excellent candidates for population genomic studies because of their large population size, continuous distribution, and diversity of habitat, few genomic resources are currently available for this species. Here, we sequence and assemble the mule deer genome into a highly contiguous chromosome-length assembly for use in future research using long-read sequencing and Hi-C. We also provide a genome annotation and compare demographic histories of the mule deer and white-tail deer using PSMC. We expect this assembly to be a valuable resource in the continued study and conservation of mule deer.

## Data Description

### Background and Context

The mule deer (*Odocoileus hemionus*) is a mid-sized ruminant [50-90 kg; Figure 1; 1, 2], ranging from the Yukon Territory in Canada to Central Mexico. Mule deer can be found in boreal forests, high and low elevation desert shrublands, subalpine forests, woodlands, prairies, and a variety of other habitats with subspecies and types frequently inhabiting different habitats [3]. They belong to the Cervidae family, one of the most speciose families in the mammal suborder Ruminantia [4]. Eleven subspecies of mule deer have been recognized but are grouped into two morphologically distinct types, mule deer (*O. h. hemionus, fulginatus, californicus, inyoensis, eremicus, crooki, peninsulae, sheldoni,* and *cerrosensis*) and black-tailed deer *[O.h. columbianus*, and *sitkensis*; 5]. While the two types are well-supported by morphological and DNA evidence, little divergence has been observed among the subspecies within each type [6, 7]. This is likely due to large population sizes and the frequency of long-distance dispersal by individual deer maintaining gene flow among populations [8, 9].

**Figure 1.**
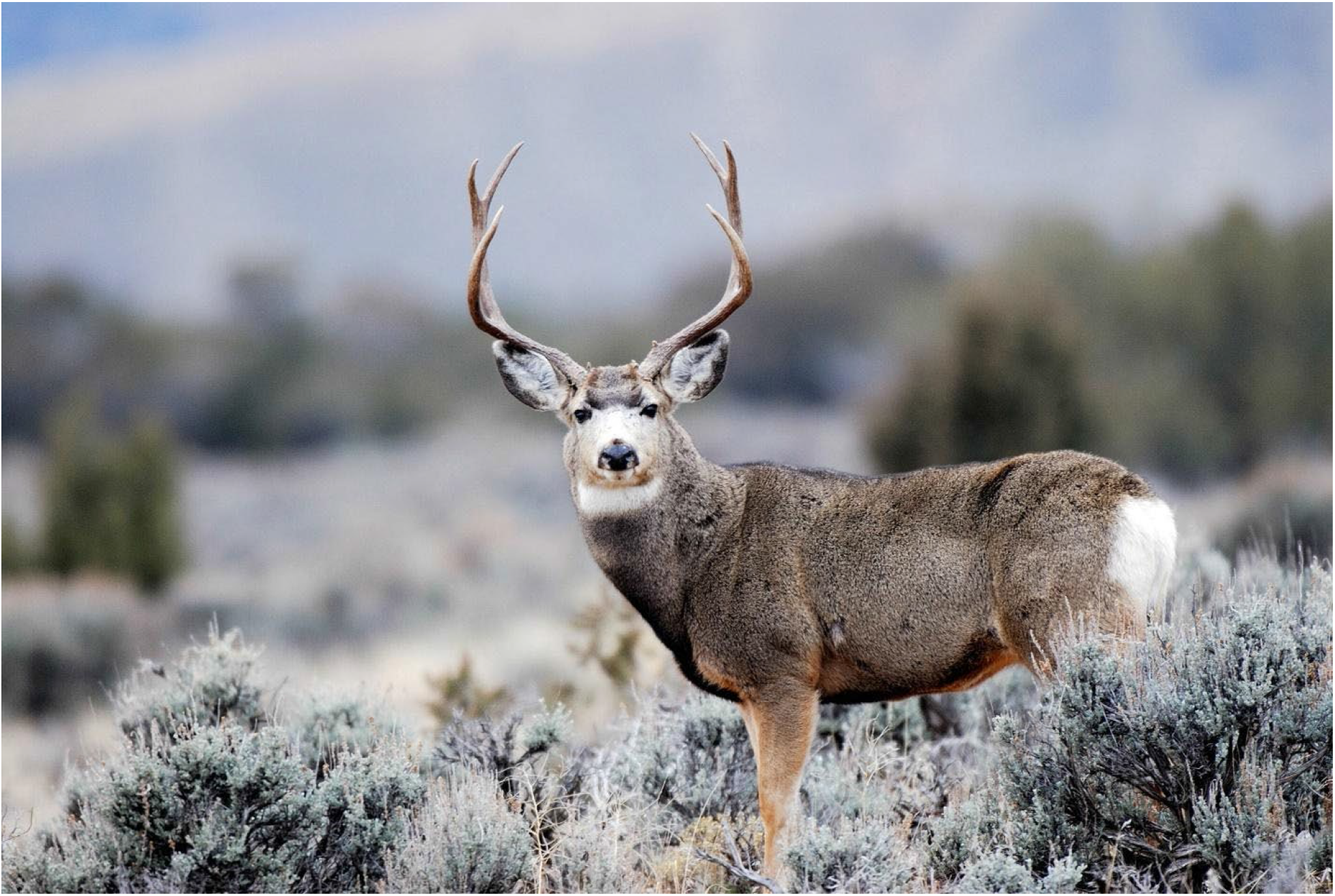
Photograph of *Odocoileus hemionus*. Photo courtesy of the Utah Division of Wildlife Resources.

**Figure 2:**
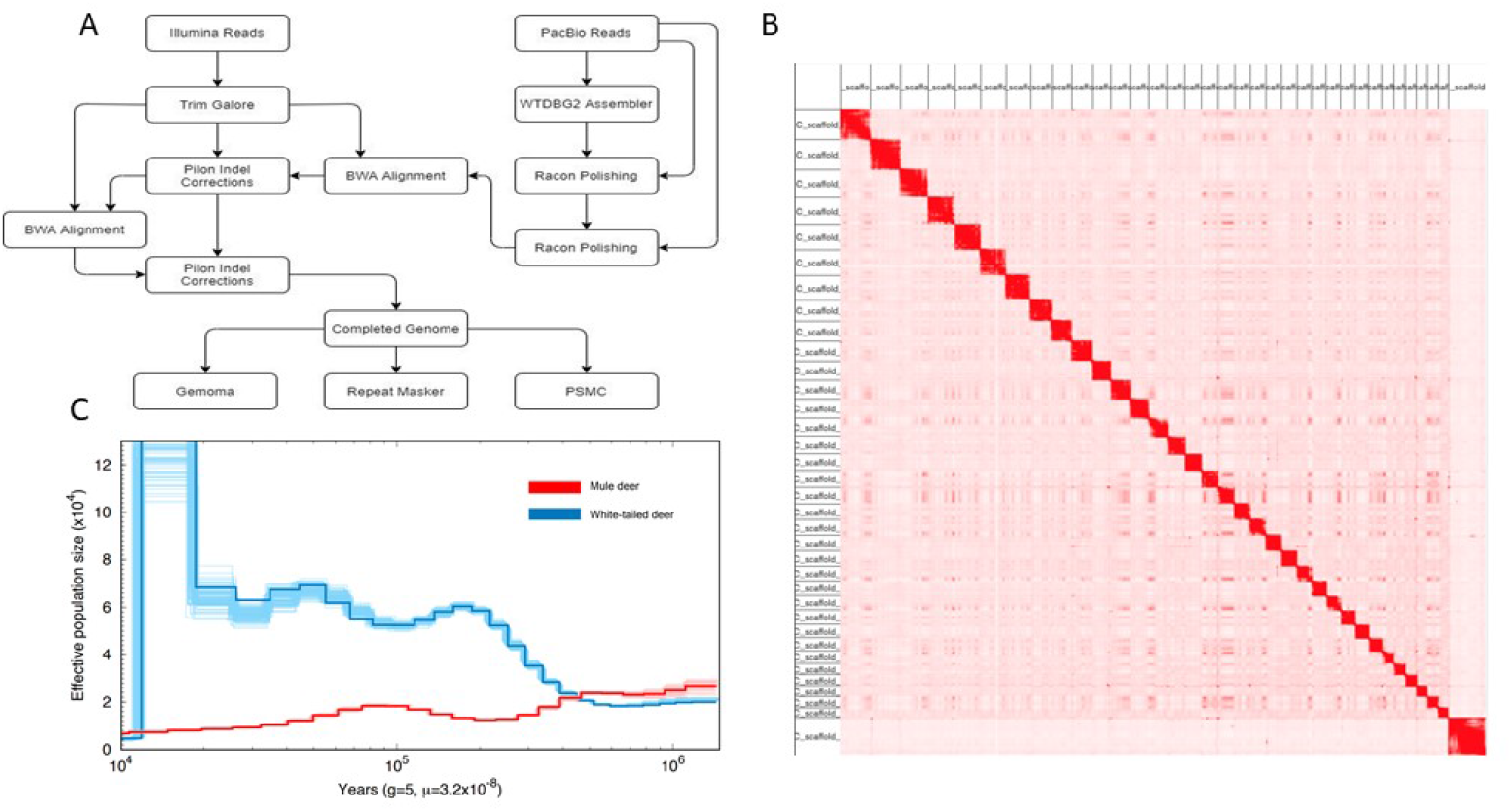
**A.** Summary flow chart of software used for the genome assembly and annotation of *Odocoileus hemionus*.**B.** Hi-C contact map of the 35 chromosome-length scaffolds. 93.45% of the genome is held in these chromosome-length scaffolds. **C.** Demographic histories estimated with PSMC for *Odocoileus virginianus* and *O. hemionus*

Characteristics such as large population size, diversity of habitat and capacity for long distance dispersal make mule deer a good candidate species for genomic study [10–12]. However, limited genomic resources are available. Currently, genetic resources available for *Odocoileus spp*. are limited to a variety of microsatellite loci [13–15] and molecular resources gleaned from the bovine genome [16–18]. Recently, Russell et al. published the first draft whole genome sequence assembly and a species diagnostic SNP panel specifically for mule deer [19]. However, this assembly was based on low-coverage short-read sequencing (Illumina) and was assembled using a reference-based approach, limiting identification of large structural variants.

Here, we report a high-quality, chromosome-length reference genome for mule deer assembled from a combination of long-read (PacBio) and short-read (Illumina) sequence data and scaffolded using Hi-C. Our goal was to develop whole genome resources that will aid in better understanding questions related to mating systems, parentage assignment, relatedness, estimation of demographic parameters, population genetic analysis, and assessment of population viability [20]. We also provide an annotation and estimate demographic histories of both the white-tail and mule deer using the Pairwise Sequentially Markovian Coalescent (PSMC) model. We discuss how this new genome assembly can be applied to conservation and management of mule deer.

## Methods

### Sample collection and DNA preparation

A tongue biopsy was collected within 2 hours postmortem from a single female mule deer that was removed for depredation purposes from Woodland Hills, Utah (40°00’ N 111° 38’ W). The biopsy was immediately stored on ice and frozen at −80° Celsius within 12 hours of collection. The sample remained frozen at −80° Celsius until DNA extraction and sequencing were performed. Genomic DNA was extracted from the tongue tissue after treating the tissue with proteinase K using a Qiagen Genomic Tip Kit for High Molecular Weight DNA following Qiagen’s extraction protocol (Qiagen, Valencia, CA, USA). After extraction, the DNA was visualized with pulsed-field gel electrophoresis to evaluate whether the DNA was of sufficient length for single-molecule real-time (SMRT) sequencing using the PacBio Sequel II sequencing instrument [21].

### Sequencing and Assembly

The DNA extractions were successful on the first attempt and the pulsed-field gel showed sufficient DNA length, with a band above 50 kbp. The extracted DNA was sheared to 65 kbp and then size-selected for fragments greater than 32 kbp using a Sage Science BluePippin. The size-selected DNA was prepared into a PacBio library using the SMRTbell® Express Template Preparation Kit 2.0 (PacBio, USA), then sequenced across two PacBio Sequel II 8M SMRT cells (PN:101-389-001). Each run was performed at the Brigham Young University DNA Sequencing Center (Provo, Utah).

Extracted DNA was also prepared into a paired-end Illumina library with a fragment size of 500 bp. The library was prepared using the NEBNext® Ultra™ II DNA Library Prep Kit for Illumina, and the manufacturer’s instructions were followed as outlined in the kit manual (New England BioLabs, Inc., USA). The library was sequenced across two Illumina HiSeq 2500 lanes with 2×150 bp paired-end sequencing at the Brigham Young University DNA Sequencing Center (Provo, Utah).

We converted the PacBio subreads BAM file to FASTQ using Samtools v.1.9 [22] and generated a first assembly using WTDBG2 v.2.5-1 with the command parameters “-x sq -g 2.3G -t 80 -L5000.” [23]. Reads shorter than 5000 bp were removed and not used in the assembly using the “-L5000” parameter in WTDBG2. The approximate genome size was estimated using the genome length of the previous mule deer genome and was set to 2.3Gbp. The consensus sequence was derived using the command “wtpoa-cns -t 80 -i” [24].

We recovered 239 giga-bases of PacBio subread data (~90x coverage) from the two PacBio Sequel II SMRT cells. The first SMRT cell generated 114.19 Gbp of subread data with a mean polymerase read length of 23,861 bp and a read n50 of 31,007 bp. The second SMRT cell generated 125.82 Gbp of subread data with a mean polymerase read length of 29,002 bp and a read n50 of 46,596 bp. The Illumina sequencing run yielded ~690 million reads equaling 87.2 Gbp of raw sequence data.

### Genome Polishing

We performed an initial error correction step by remapping the PacBio long reads back to the WTDBG2 contig assembly sequence using Minimap2 v.2.17-r941 “ -ax map-pb -t 40” and sorting, indexing, and converting the alignment file with the command “sort -o -T reads.tmp” and “index reads.sorted.bam” in Samtools v.1.9 into BAM format. We performed two rounds of Racon error correction using “-u -t 80” parameters with the PacBio reads, with a separate alignment file created for each run.

We conducted genome polishing with high fidelity short-read data by first mapping Illumina reads to the Racon corrected consensus assembly. We first trimmed adapters from the Illumina sequences using Trim Galore v.0.6.4. We then mapped Illumina reads to the Racon corrected assembly using BWA v.0.7.17-r1188 and sorted and indexed the alignment file with Samtools v.1.9. We used Pilon v.1.23 to correct indel errors using “--vcf --tracks --fix indels -- diploid” parameters. We then ran a second round of indel correction by repeating the steps above on the output from the first round of Pilon.

We generated assembly statistics using the assembly_stats script [25] and used BUSCO v. 5.2.2 used to evaluate the recovery of universal single copy orthologs using the mammalia_odb9 ortholog set [26]. The assembled mule deer genome has a total length of 2.61 Gbp with a GC content of 41.8% and a contig N50 of 28.6 Mbp (Table 1) with a longest contig of roughly 96.5 Mbs.

**Table 1.** Metrics of *O. hemionus* genome assembly.

| Statistic | Contig Statistics | Scaffold Statistics |
| --- | --- | --- |
| L10 | 3 | 2 |
| L20 | 7 | 4 |
| L30 | 11 | 7 |
| L40 | 17 | 10 |
| L50 | 25 | 13 |
| N10 | 70,175,673 | 112,787,451 |
| N20 | 61,053,151 | 102,606,451 |
| N30 | 53,514,773 | 91,763,312 |
| N40 | 41,740,608 | 76,003,156 |
| N50 | 28,570,879 | 72,140,960 |
| Longest | 96,465,089 | 139,398,775 |
| Mean | 432,466 | 473,563 |
| Median | 23,026 | 20,939 |
| Sequence Count | 6,033 | 5,510 |
| Shortest | 1,000 | 1,000 |
| GC Content | 41.8 | 41.8 |
| Total Base Pairs | 2,609,071,524 | 2,609,333,024 |

### Chromosome-length Scaffolding

High-throughput chromosome conformation capture (Hi-C) was performed to provide chromosome-length scaffolding for the consensus genome (Figure 3). Data generation and Hi-C scaffolding was performed by the DNA Zoo Consortium (https://www.dnazoo.org). In brief, in situ Hi-C data [27] was aligned to a draft genome assembly using the Juicer pipeline [28]. The 3D-DNA pipeline [29] was used to error-correct, anchor, order and orient the pieces in the draft assembly, producing a candidate assembly. The candidate assembly was manually reviewed and polished using Juicebox Assembly Tools aka JBAT [28, 30]. Interactive contact maps visualized using Juicer.js [31] for before and after the Hi-C scaffolding are available at https://www.dnazoo.org/assemblies/Odocoileus_hemionus.

**Figure 3:**
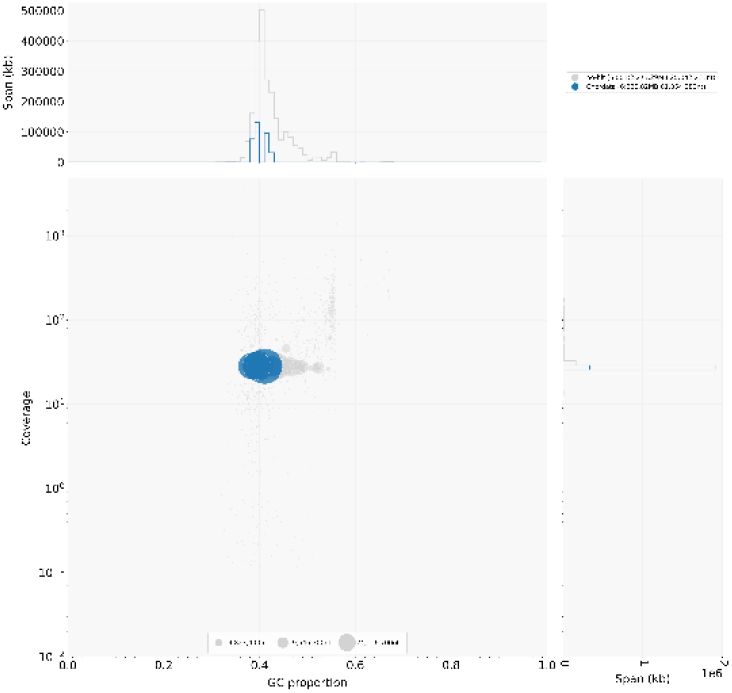
Blobplot of the mule deer genome assembly. All contigs are assigned to Chordata.

The Hi-C scaffolding placed 93.45% of the total base pairs in the assembly into chromosomes. We successfully identified 94.5% of BUSCO genes in the assembly, with 91.2% single copy and 3.3% of duplicated BUSCOs, comparable to other recently published cervid genomes [Table 2; 32].

**Table 2.** BUSCO v. 5.2.2 statistics.

| BUSCO Statistic | Number Identified | Percent Identified |
| --- | --- | --- |
| Complete | 8719 | 94.5% |
| Complete and Single-Copy | 8412 | 91.2% |
| Complete and Duplicated | 307 | 3.3% |
| Fragmented | 104 | 1.1% |
| Missing | 403 | 4.4% |
| Total | 9226 | 100% |

### Genome annotation

RepeatMasker (http://www.repeatmasker.org/) was used with the NCBI engine to estimate the overall repeat content of the genome [33]. Repeat databases were built using RepeatModeler v.2.0.1 with parameters “BuildDatabase -name -engine ncbi && RepeatModeler -engine ncbi -pa 8 -database”. RepeatMasker v.4.1.1 was used to identify repeats using the parameters “-pa 16 - gff -nolow -lib” (Table 3).

**Table 3.** RepeatMasker Annotation.

| Annotation Type | Percentage |
| --- | --- |
| Retroelements | 31.72% |
| DNA transposons | 1.38% |
| Rolling-circles | 1.62% |
| Interspersed Repeats | 37.86% |
| Small RNA | 0.80% |
| Satellites | 0.95% |
| Simple Repeats | 0.04% |
| Unclassified | 4.76% |

We performed homology-based gene prediction using Gene Model Mapper (GeMoMa) v.1.6.4 with the existing *Odocoileus virginianus* [34] genome annotation used as a reference; the following command was used “GeMoMa -Xmx50G GeMoMaPipeline threads=40 outdir=annotation_out GeMoMa.Score=ReAlign AnnotationFinalizer.r=NO o=true t=mule_deer.fa i=white_tail a=GCF_002102435.1_Ovir.te_1.0_genomic.gff g=GCF_002102435.1_Ovir.te_1.0_genomic.fna”. The GeMoMa annotation predicted 21,983 full-length proteins.

Blobtools v1.1.1 was used to evaluate the assembled genome for possible contamination. Blast v.2.9.0 was used to identify any possible contamination using the command “blastn -task megablast -outfmt ‘6 qseqid staxids bitscore std’ -max_target_seqs 1 -max_hsps 1 -num_threads 16 -evalue 1e-25”. A blobplot was created for visualization using the “create” function of blobtools. The blobplot revealed no evidence of contamination in the genomes (Figure 4).

### Historical demography

We used the Pairwise Sequentially Markovian Coalescent (PSMC) v.0.6.5-r67 to estimate the demographic history of the mule deer [35]. We re-aligned Illumina reads to the final assembly with BWA and sorted and indexed the alignment file in Samtools v.1.9. We used mpileup and bcftools to call heterozygous sites using the command “samtools mpileup -C50 -uf” and “bcftools call -c” respectively. Additionally, Bcftools v.1.11 was used with the vcfutils.pl utility and the following parameters “vcf2fq -d 10 -D 90”. We then used PSMC v.0.6.5-r67 to generate the demography history. We first created a psmcfa file with the following command “fq2psmcfa -q20”. The psmcfa file was split using “splitfa” function of PSMC. A PSMC was created using the command “psmc -N25 -t15 -r5 -p ‘4+25*2+4+6’. Bootstraps were created from the split psmcfa file using the command “seq 100 | xargs -i echo psmc -N25 -t15 -r5 -b -p “4+25*2+4+6” \ -o round-[36].psmc $splitpsmcfa | sh” The initial PSMC and bootstraps were then merged and visualized with psmc_plot.pl using the command “psmc_plot.pl -pY20 -g5 -u 3.22e-8”. To compare demographic histories with the other most common North American deer species, the white-tailed deer (*Odocoileus virginianus*), we followed the same process described above. We downloaded the *O. virginianus* assembly from NCBI (accession: NC_015247) and downloaded the raw Illumina reads from the sequence read archive (SRA) using the fastq-dump, utility within SRAtoolkit v.2.10.9, with the following parameters “fastq-dump --gzip --skip-technical --readids --read-filter pass --dumpbase --split-e –clip”. Because fastq-dump alters read names, individual read names were corrected to match in both the forward and reverse fastq files by removing “.1” from the end of the forward reverse identifier and “.2” from the end of the reverse sequence identifier.

We used a PSMC analysis to compare historic population trends of *O. hemionus* and *O. virginianus*. In comparing the PSMC analysis, we observe that *O. hemionus* and *O. virginianus* have divergent demographic histories. As effective population size for *O. hemionus* increases, the effective population size of *O. virginianus* appears to decrease, and vice versa. The effective population size of *O. hemionus* has been in a constant decline since the most recent glacial period roughly 500,000 years ago. Two possible explanations for this decline may include overall population decline or population fragmentation. This pattern is divergent from *O. virginianus* which has shown increases in effective population size since the same time period. While both deer species inhabit the same continent, and even possess some overlapping habitat, it appears that the species react differently to environmental changes (Figure 4).

### Re-use potential

Our high-quality draft genome of the mule deer represents an advance in available genomic data for the *Odocoileus* genera. With a total length of 2.6 Gb and a contig N-50 of 28.6Mbs, this chromosomal-level *de novo* assembly can serve as a base for future conservation and genomics research. Due to the importance of deer on both the ecosystem level and to local economies, a continued effort to conserve these populations is vital [37]. Our hope is that this genomic resource will further our understanding of *O. hemionus*, and subsequently lead to more effective management of the species, including insights into the impact of anthropogenic barriers on gene flow, the possibility of species divergence in isolated populations, and the presence of multiple paternity [38, 39].

## Data Availability

The *Odocoileus hemionus* genome and raw reads are publicly accessible through NCBI. The genome data is available via BioProject ID: PRJNA752226. The raw Hi-C data is available via PRJNA512907. The data sets supporting the results of this article are available at the following link https://byu.box.com/v/mule-deer-genome

## Declarations

### List of abbreviations

bp: base pair
BUSCO: Benchmarking Universal Single-Copy Orthologs
Gb: gigabase
Hi-C: high-throughput chromosome conformation capture
kb: kilobase
Mb: megabase
NCBI: National Center for Biotechnology Information
PSMC: Pairwise Sequentially Markovian Coalescent
SMRT: single-molecule real-time

## Author’s Contributions

S.L., P.B.F, R.T.L, and B.R.M. designed this project. A.M.T. prepared the samples. A.M.T., P.B.F. performed the analyses. R.K., D.W., O.D. and E.L.A. generated the Hi-C data and performed the scaffolding. S.L., A.M.T., T.A.H., P.B.F., B.R.M. and R.T.L. wrote and revised the manuscript.

## Acknowledgements

We also thank Seth B. Wilson and Ethan Tolman for assistance with our GenBank upload and PSMC. All computation was performed on the Fulton Supercomputer (Brigham Young University, Provo, Utah, USA). Hi-C data for the mule deer were created by the DNA Zoo Consortium (https://www.dnazoo.org). DNA Zoo sequencing effort is supported by Illumina, Inc.; IBM; and the Pawsey Supercomputiong Center, E.L.A. was supported by an NSF Physics Frontiers Center Award (PHY1427654), the Welch Foundation (Q-1866), a USDA Agriculture and Food Research Initiative Grant (2017-05741), and an NIH Encyclopedia of DNA Elements Mapping Center Award (UM1HG009375).

